# Missing steps of mitochondrial translation initiation identified in plants

**DOI:** 10.64898/2025.12.30.697032

**Authors:** Vasileios Skaltsogiannis, Heddy Soufari, Philippe Giegé

## Abstract

Translation initiation, the first step of the translation cycle, involves initiation factors (mtIF) 2 and 3 in mitochondria. mtIF3 release from the ribosomal small subunit was believed to be a prerequisite for the recruitment of the initiator tRNA. Here we use cryogenic electron microscopy to characterize plant mitochondria pre-initiation complexes and to reveal how plant mtIF3 binds the initiator tRNA and facilitates its accommodation in the decoding center.

---

Translation is the final step of gene expression that converts genetic information encoded by mRNAs into proteins. Mitochondrial translation has attracted considerable interest in the recent years as it combines bacterial features inherited from the prokaryote ancestor of mitochondria with several eukaryote specific traits ^1–4^. As part of the translation cycle, the initiation step utilizes the initiation factors 2 and 3 in both bacteria and mitochondria ^5,6^. Mitochondrial initiation factors (mtIF) 2 and 3 resemble the bacterial ones, although they are defined by the occurrence of eukaryote specific domains ^7–10^. While IF2 function for the positioning of initiator formylated tRNA-Met (fMet-tRNA^iMet^) appears to be comparatively conserved in bacteria and mitochondria, the function of mtIF3 has been subjected to debate^11^. In bacteria, IF3 interferes with the ribosome small subunit (SSU) and large subunit (LSU) association. It guarantees that fMet-tRNA^iMet^ is accurately chosen instead of elongator aa-tRNAs and it contributes to rejecting mRNAs that possess suboptimal translation initiation sequences ^12,13^. In contrast, in animal mitochondria, mtIF3 was found to coordinate the late step of SSU assembly as it binds to the ribosome before the mitoribosomal protein mS37. At this stage IF3 interacts with the assembly factor RBFA. The latter is then replaced by mS37 which is the last protein incorporated to complete the assembly of the SSU ^14^. mtIF3 also controls the formation of the pre-initiation complex (mtPIC). For this, mtIF3 forms extensive contacts with the mtSSU platform, preventing the premature association of mtPIC with the LSU and amino-acylated initiator tRNA. The availability of *in vitro* reconstitution and structural analyses of the mammalian mtPICs have enabled to propose a model for mitochondrial translation initiation. In this model, mRNA loading and fMet-tRNA^iMet^ recruitment take place at the same time as mtSSU and mtLSU join to form the monosome and strictly depend on prior release of mtIF3 ^8,14^. Here, we bring evidence that allows to reconsider this model. We use cryogenic electron microscopy (cryo-EM) to reveal a new function of mtIF3. Results show that plant mtIF3 binds fMet-tRNA^iMet^ at the level of the mtSSU, with the N-terminal domain of mtIF3 helping tRNA accommodation to the decoding center. We also observe how mRNA is recruited before the release of mtIF3. Whether these steps of translation initiation using a novel initiator tRNA binding activity of mtIF3 are specific to plants or whether they were missed from previous studies using different eukaryote models is discussed.

To get insights into translation initiation in plant mitochondria, orthologs of bacterial IF2 and IF3 were identified in Arabidopsis mitochondria as being At4g11160 (mtIF2) and At1g34360 (mtIF3) respectively. The localization of these proteins was confirmed to be mitochondrial by confocal microscopy (Supplementary Fig. 1a). Arabidopsis mtIF2 resembles both bacterial and human mtIF2, although it does not contain the 37 amino acids domain inserted in human mtIF2 that has taken the function of IF1 ^7,15,16^. Likewise, Arabidopsis mtIF3 is structurally similar to the bacterial and human mtIF3. Nevertheless, Arabidopsis mtIF3 has a C-terminal additional domain that doubles the size of the protein (Supplementary Fig. 2). The analysis of knock-out mutants performed here showed that both Arabidopsis mtIF2 and 3 are essential proteins. In human, mtIF2 is also essential, however, mtIF3 is dispensable for mitochondrial translation in cultured human cells ^17^, yet heart and skeletal muscle depletion of mtIF3 causes cardiomyopathy in mice and leads to death beyond 20 weeks of age ^18^. In the context of translation, mtIF3 has never been detected together with the initiator tRNA or the mRNA in any of the described pre-initiation complexes, suggesting that it is not required for monosome formation ^8,14^. Still, mtIF3 is essential for the translation of second ORFs in bicistronic transcripts, although its role in the translation of leaderless transcripts is debated ^19,20^. In order to assess the mode of action of plant mtIF2 and 3 and to decipher their role in translation initiation, Arabidopsis mtIF2 and 3 were expressed in a cell-free protein synthesis system using plant extracts and purified to homogeneity (Supplementary Fig. 1b). Likewise, Arabidopsis mitochondrial initiator tRNA was transcribed, amino-acylated and formylated *in vitro* to obtain fMet-tRNA^iMet^, as confirmed by mass spectrometry analysis (Supplementary Fig. 3).

As a next step, plant mitochondrial translation pre-initiation complexes were reconstituted *in vitro* by incubating different combinations of plant mitoribosome subunits, purified as described previously ^21,22^, Arabidopsis mtIF2, mtIF3, fMet-tRNA^iMet^, and a 51-nt transcript representing rps3 mRNA. Three reconstitution reactions were prepared: the first contained mtSSU, mtIF3, and rps3 mRNA; the second consisted of mtSSU, mtIF3, rps3 mRNA, mtF2, and fMet-tRNA^iMet^, whereas the third reaction included the components of the second with the addition of mtLSU. Two separate datasets were collected per reaction resulting in the four structures presented here: (1) a complex of the mtSSU with mtIF3, which we consider the earliest step toward the complete initiation complex (IC) and termed mtPIC-1; (2) a complex of mtSSU-mtIF3-mRNA, identified as mtPIC-2; (3) a complex of mtSSU-mtIF3-mRNA-tRNA, identified as mtPIC-3; and finally (4) a complex of mtSSU-mtLSU-tRNA that represents the initiation complex (mtIC*) after the dissociation of all auxiliary factors (Fig. 1).

**Figure 1:**
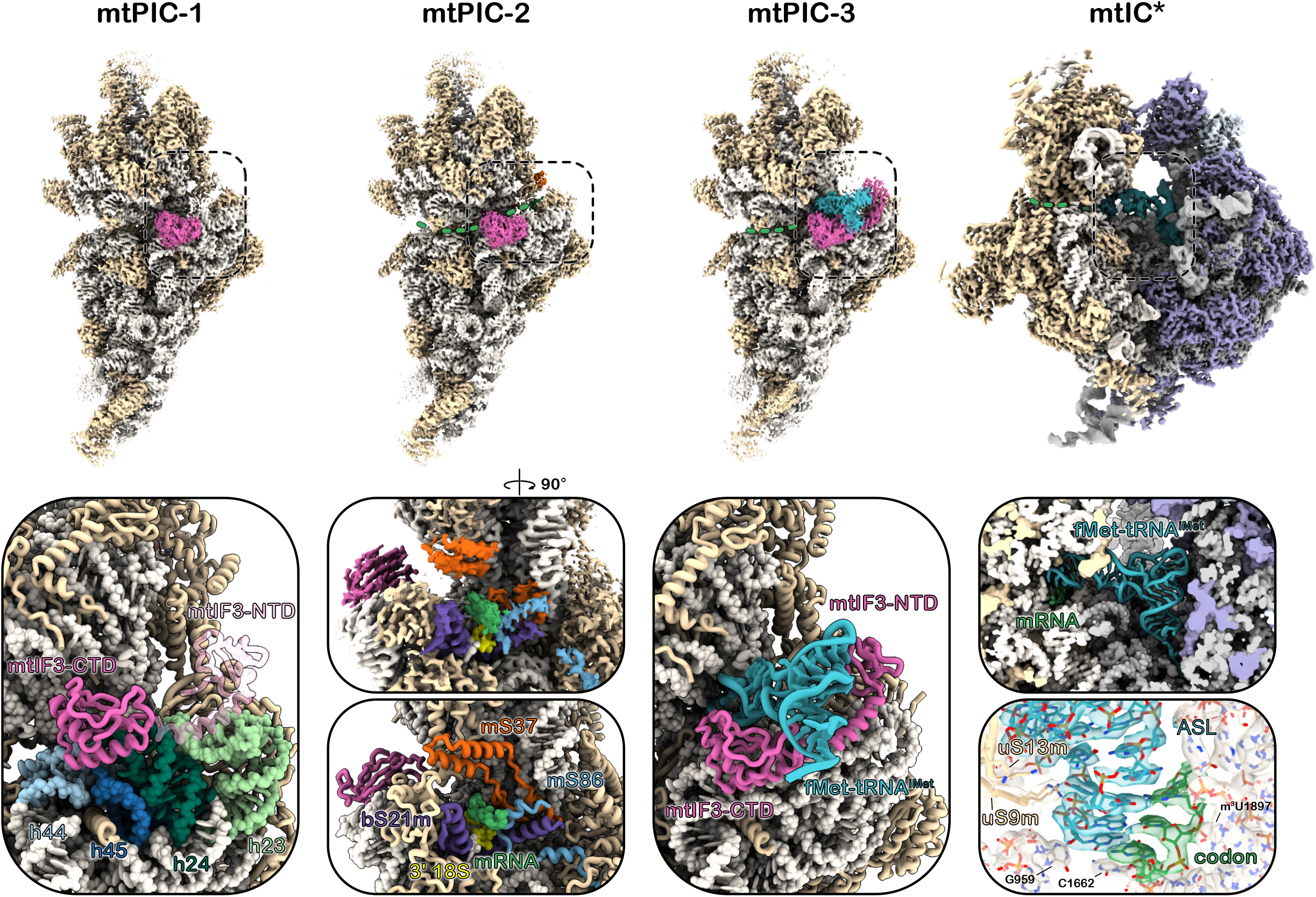
Plant mitochondrial pre-initiation complexes. Cryo-EM maps of mitochondrial pre-initiation complexes mtPIC-1, mtPIC-2, mtPIC-3 and initiation complex mtIC* (top) with their atomic models in close-up view (below). **mtPIC-1** mtIF3-CTD resides on top of h44, h45 and h24 whereas the most flexible NTD (transparent) is positioned in the vicinity of the platform yet remains unresolved at high resolution. **mtPIC-2** close-up view of the mRNA channel exit site with both density and model depicted. Density attributed to mRNA is resolved as it is stabilized by mS37, mS86, bS21m and 3’ 18S. **mtPIC-3** mtIF3 recruits the initiator tRNA to the decoding center. **mtIC\*** the initiation complex following the dissociation of auxiliary factors. zoomed-in view of the initiator tRNA and the codon-anticodon interaction. mtIF3, initiator tRNA, mRNA, rRNA elements and r-proteins are consistently coloured across panels.

Analysis of the first reaction led to the identification of two pre-initiation complexes, mtPIC-1 and mtPIC-2 resolved at 2.28 Å and 2.55 Å resolution, respectively. mtPIC-1 represented 33% of the particles, whereas mtPIC2 the 15%, with the overall number of mtSSU particles accommodating mtIF3 reaching 70% of their total number. In mtPIC-1, mtIF3 binds to the decoding center of the mtSSU in a conformation where its C-terminal domain (CTD) docks onto the rRNA helices h44 and h24 and its linker spans across h23 toward the mtSSU platform allowing the N-terminal domain (NTD) to reach the mitochondrial conserved r-protein mS37. This specific conformation of mtIF3 prevents the association of mtLSU, mainly via the placement of its CTD that would clash with h69 of the mtLSU upon subunit joining (Fig. 1) (Supplementary Fig. 4). In mtPIC-2, the initiation factor mtIF3 retains its position and conformation on the mtSSU interface whereas the mRNA channel is occupied by the rps3 mRNA (Fig. 1) (Supplementary Fig. 4). During the formation of these early pre-initiation complexes, the NTD exhibits high flexibility and as a result its position is only traceable in low contour threshold. Analysis of the second reconstitution reaction revealed a subpopulation of 5.7 % of the total particles that represents the mtSSU-mRNA-mtIF3-tRNA complex, resolved at 3.05 Å resolution and assigned as mtPIC-3 (Fig. 1) (Supplementary Fig. 5). In this state, mtIF3 holds a central role in the recruitment of the initiator tRNA and its accommodation to the P site (Fig. 2a-e). The mtIF3-NTD interacts with the elbow of the initiator tRNA (Fig. 2e), while the linker encircles the tRNA reaching its concave side where the CTD resides. The CTD then funnels the tRNA anticodon stem loop (ASL) into the P site by forming a barrier with the residues 196-201 and 233-247. Moreover, the loop ^204^DKDKHK^209^ probes the tRNA ASL to facilitate the codon-anticodon interaction, that is further reinforced by the rRNA residue C1662 which secures the anticodon position by stacking with C35. Nevertheless, the codon-anticodon interaction takes place only partially, as only one out of the three base pairings is unambiguously resolved (Fig. 2c). An additional function of the CTD is the stabilization of the tRNA acceptor arm. This is achieved by the mitochondria-specific C-terminal extension (CTE) (residues 255-260) which extends beneath the acceptor arm providing a steady support (Fig. 2d). Despite the establishment of the codon-anticodon recognition, the overall positioning of the tRNA differs significantly from the final one, as adopted during the elongation state (Fig. 2f). This can be obtained either via the relocation of the CTD, which currently sterically precludes the tRNA from taking its final configuration or the dissociation of mtIF3 entirely. In our third reconstitution reaction we added the mtLSU in order to trigger the formation of the initiation complex. The majority of the mitoribosomal subunits (75% of the particles) assembled into the full ribosome, resolved at 2.92 Å resolution. The formed monosome accommodated the rps3 mRNA along the mRNA channel and harboured the initiator tRNA in the P site, hallmarks of the initiation complex (representing 6% of the monosome particles). In contrast to mtPIC-3, the initiator tRNA here adopts its final configuration, with the acceptor stem and elbow being perpendicular to the ASL. Since all auxiliary factors were absent, we named this step mtIC* (Fig. 1) (Supplementary Fig. 6).

**Figure 2:**
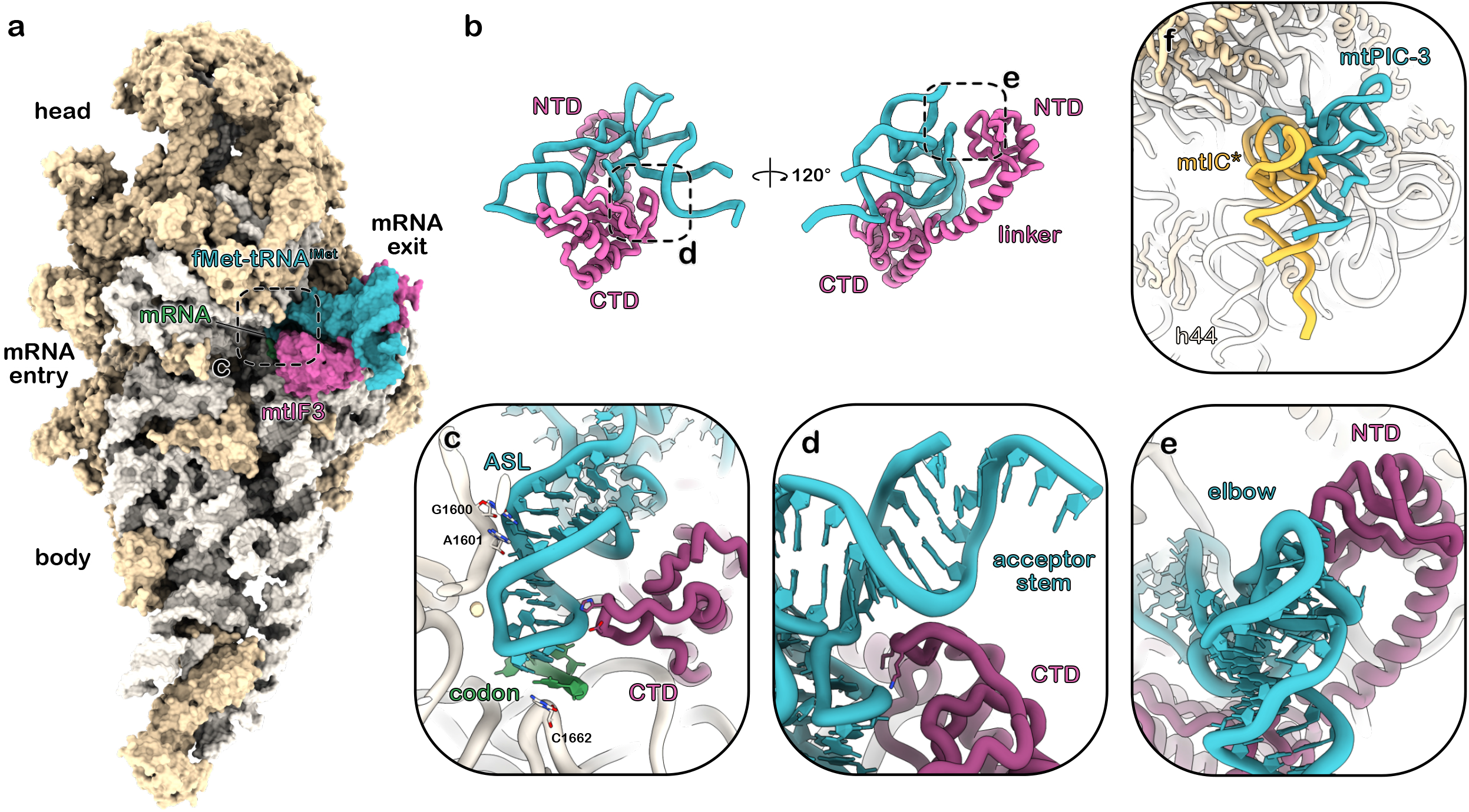
The plant mitochondrial initiation factor 3 recruits the initiator tRNA. **a** Composite map of the mitochondrial pre-initiation complex 3 displaying mtIF3 in association with the initiator tRNA at the decoding center of the mtSSU. **b** Close-up view of the atomic model of mtIF3-tRNA complex, shown from two perspectives. Dashed boxes highlight regions presented at higher detail. **c** mtIF3-CTD probes the initiator tRNA ASL promoting the codon-anticodon recognition, and **d** stabilizes the acceptor stem, **e** whereas the NTD interacts with the tRNA elbow. **f** Comparison of the two discrete configurations of the initiator tRNA between mtPIC-3 and mtIC* (yellow).

Collectively, an *in vitro* reconstitution approach allowed us to capture four distinct complexes of the plant mitochondrial translation initiation pathway. Three correspond to the pre-initiation phase (mtPIC-1 to mtPIC-3) that progressively lead to a translation-ready initiation complex (mtIC*) (Fig. 2). The mtPIC-1 identified here is equivalent to the early pre-initiation complex mtSSU-mtIF3 that has been described in human mitochondria ^23^, suggesting that the early steps of translation initiation in human and plant mitochondria converge. Nevertheless, the following pre-initiation steps identified here, are in striking difference with those observed in mitochondria of humans or other eukaryotes. We report two additional pre-initiation complexes featuring mtIF3, one in association with the translating mRNA (mtPIC-2) and the other in association with both mRNA and initiator tRNA (mtPIC-3). This constitutes the first observation of mtIF3 in complex with either the mRNA or the initiator tRNA, whereas prior to this work the dissociation of mtIF3 was generally recognized as a requirement for the subsequent accommodation of the initiator tRNA. Thus, plant mitochondria appear to follow a more bacterial-like sequence of events along their translation initiation pathway ^12^. Moreover, it becomes apparent that plant mitochondria have reserved an orchestrating role for mtIF3 during translation initiation to accommodate the initiator tRNA in the decoding center and properly orient it for start codon recognition. In contrast, the current model, derived from structural analyses mainly in mammalian mitochondria, chronically places the accommodation of the mRNA and the initiator RNA with the recruitment of mtIF2 upon monosome assembly ^7,14^. Nevertheless, it remains unanswered whether the mRNA and the initiator tRNA can be recruited to the mtSSU prior to the mtLSU joining or if monosome formation is a prerequisite. Our findings have enabled us to describe a sequence of discrete steps that mitoribosomes undergo during translation initiation in plant mitochondria. These steps are presented in the following model: in mtPIC-1, mtIF3 is first recruited on the mtSSU, presumably replacing the late assembly factor RsgA ^21^, followed by the accommodation of the mRNA along the mRNA channel, mtPIC-2. *In vivo*, the mRNA delivery is possibly directed by translation trans-factors (TTF), similar to the LRPPRC-SLIRP complex in human ^24^ or translational activators in yeast ^25^. In mtPIC-3, the initiator tRNA is recruited to the mtSSU-mtIF3-mRNA complex. mtIF3 facilitates the accommodation of the initiator tRNA in the P site and promotes start codon recognition. Although, absent from our data, the next step, mtPIC-4, should require the association of mtIF2, to reorient the tRNA position in its final configuration, whereas mtIF3 dissociates. The inability to detect mtPIC-4 most probably reflects a very transient interaction of mtIF2 with mitoribosome subunits in plants. Finally, the initiation complex (mtIC) is formed upon mtLSU joining, followed by the departure of mtIF2 leading to a translation-ready initiation complex (mtIC*) (Fig. 3). Considering that the key players described above are conserved across eukaryotes, we suggest this model might be applicable beyond plants. In fact, despite, the divergence in amino acid sequence between plant and human mtIF3, structurally the two proteins adopt a similar conformation, retaining the two conserved domains crucial for the interaction with the initiator tRNA (Supplementary Fig. 2b). Thus, an analogous role for the human mtIF3 that will restore the presence of the mRNA and initiator tRNA in pre-initiation complexes prior to monosome formation, cannot be excluded. Future studies will show whether plants have followed a distinct evolutionary route or whether the steps of mitochondrial translation initiation involving mtIF2 and 3 are universal across eukaryotes.

**Figure 3:**
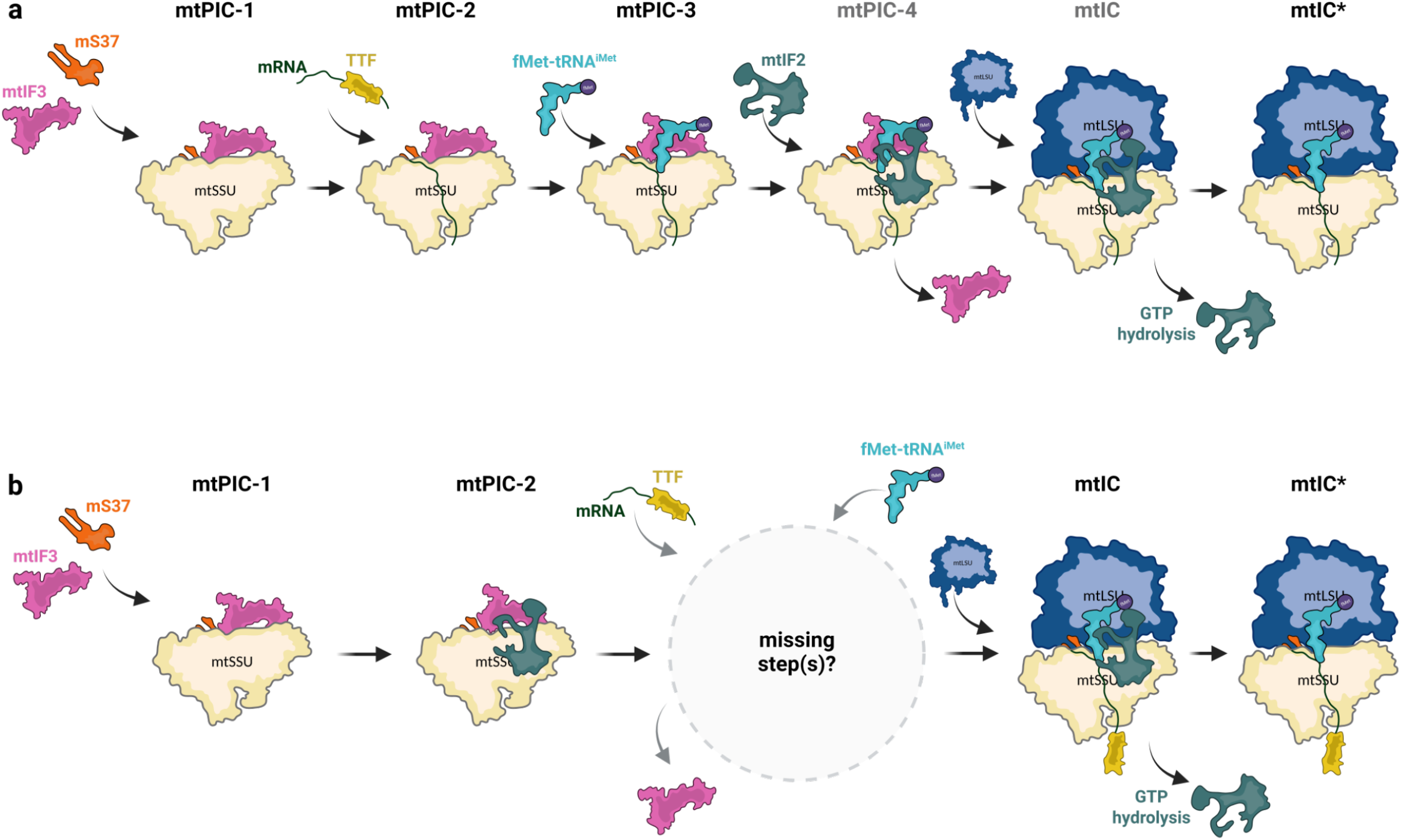
Schematic representation of the sequence of events leading to translation initiation in mitochondria. **a** model deduced from the structures of plant mitochondria initiation complexes presented in Fig. 1. This representation comes as an alternative to the model based on studies performed in human mitochondria shown in **b** with a schematic representation derived from Brischigliaro *et al*. ^*11*^. mtPIC are mitochondrial pre-initiation complexes, whereas mtIC are mitochondrial initiation complexes. mtSSU and LSU represent mitoribosome small and large subunits respectively. mtIF2 and 3 are mitochondrial initiation factors 2 and 3. TTF represent translation *trans*-factors required for the recruitment of mRNA to the mitoribosome, such as the LRPPRC-SLIRP complex in human, or putative PPR *trans*-factors in plants. mtPIC-4 and mtIC complexes in **a** are hypothetical. Figure created in BioRender (biorender.com).

## Methods

### Mitochondrial ribosome subunits purification

Mitochondria were purified from cauliflower as previously described ^26,27^. Mitoribosome purification was performed as previously ^21,22^. In brief, purified mitochondria were re-suspended in Monosome buffer (20 mM HEPES-KOH, pH 7.6, 75 mM KCl, 20 mM MgCl_2_, 1 mM DTT,, supplemented with proteases inhibitors (c0mplete EDTA-free)) containing 1.5% Triton X-100 to a 1.3 mg/ml concentration and incubated for 10 min at 4°C. Lysate was clarified by centrifugation at 25,000 g, 20 min at 4°C. The supernatant was loaded on a 40% sucrose cushion in Monosome buffer supplemented with 0.1% Triton X-100 and centrifuged at 235,000 g for 3h at 4°C. The crude ribosome pellet was re-suspended in Monosome buffer and loaded on a 10-30 % sucrose gradient in the same buffer and centrifuged for 15 h at 86,000 g. Fractions corresponding to full mitoribosomes were collected, pelleted and re-suspended in Dissociation buffer (20 mM HEPES-KOH, pH 7.6, 300 mM KCl, 5 mM MgCl_2_, 1 mM DTT, supplemented with proteases inhibitors (C0mplete EDTA-free)) and incubated for 2h at 4°C. The dissociated subunits were separated in a 10-30 % sucrose gradient by centrifugation for 15 h at 86,000 g. Fractions corresponding to the two mitoribosomal subunits were collected, pelleted, re-suspended in Monosome buffer, flash-frozen in liquid nitrogen and stored at -80°C for further use.

### Preparation of plant mitochondrial initiation factors 2 and 3

The plant mitochondrial initiation factors 2 and 3 were prepared using the cell-free expression system ALiCE (LenioBio). The coding sequences of mIF2 and mIF3 were fused with an N-terminal His tag, with a TEV cleavage site placed in between. The cassettes were introduced in cell-free expression system vector, pALiCE01 before they were added to the cell-free reaction. The transcription-translation reaction mix was incubated at 25 °C for 48 h. For the purification, the reaction was diluted 10 times in binding buffer (50 mM HEPES-KOH pH7.6, 500mM KCl, 5 mM MgCl_2_, 10% glycerol, 2 mM DTT, 0.1% TritonX-100, cOmplete protease inhibitor), and centrifuged at 16,000g for 10 min. The supernatant was incubated with NiNTA beads for 90 min at 4 °C. The beads were washed with binding buffer before the proteins were eluted with elution buffer (binding buffer supplemented with 250 mM Imidazole). The eluted proteins were incubated with 10 ng/μL TEV protease overnight at 4 °C to remove their tags and subsequently, they were concentrated with an Amicon Ultra 0.5 mL centrifugal filter and were flash-frozen in liquid nitrogen.

### Preparation of E. coli MetRS and FT

The coding sequences of methionyl-tRNA synthetase (MetRS) and methionyl-tRNA formyl transferase (FT) were acquired in a plasmid vector carrying a C-terminal His tag. The proteins were expressed in *E. coli* BL21 rosetta cells at 37 °C for 3 h. The enzymes were purified using their His tag with a HisTrap FF 5-mL column. Initially, the cells were lysed in lysis buffer (50 mM HEPES-KOH pH 7.6, 1 M NH_4_Cl, 10 mM MgCl_2_, 5 mM β-mercaptoethanol, 40 mM Imidazole) using a microfluidizer (LM20), before the lysates were clarified at 18,000g for 45 min at 4 °C. The clarified lysates were introduced in a HisTrap FF 5-mL column, and they were eluted with elution buffer (50 mM HEPES-KOH pH 7.6, 10 mM MgCl_2_, 5 mM β-mercaptoethanol, 500 mM Imidazole). The eluted samples were subsequently desalted and concentrated in HiPrep Desalting 26/10 before being subjected to size-exclusion chromatography on a Superdex 200 Increase 5/150 in storage buffer (50 mM HEPES-KOH pH 7.6, 100 mM KCl, 10 mM MgCl_2_, 5 mM β-mercaptoethanol).

### In vitro transcription, charging, and formylation of the plant initiator tRNA^iMet^

The sequence of the plant initiation tRNA^iMet^ was introduced in a pGEM-T vector and used as a template for PCR amplification. The T7 promoter sequence was added to the PCR product via the forward primer before it was added to a T7-guided *in vitro* transcription reaction. The *in vitro* reaction was performed using the RiboMAx T7 Large Scale System (Promega) and was set at 37 °C for 6 h according to the manufacturer’s instructions. The produced transcripts were treated with Turbo DNase at 37 °C for 1 h and they were subsequently purified with phenol/chloroform extraction and G-50 columns (GE). To induce folding of the initiation tRNA^iMet^, it was heated to 80°C for 5 min and then allowed to gradually cool to room temperature for 20 min before being placed on ice. For the aminoacylation reaction, 2 μg of folded tRNA^iMet^ was incubated in aminoacylation buffer (50 mM HEPES-KOH pH 7.6, 100 mM NaCl, 10 mM MgCl_2_, 5 mM beta-mercaptoethanol) supplemented with 2 mM L-methionine, 5 mM ATP, and 0.4 μM MetRS at 30 °C for 40 min. Then the charged Met-tRNA^iMet^ was immediately formylated for 15 min with the addition of 1 μM FT and 300 μM of the formyl donor 10-CHO-THF. The aminoacylated and formylated initiator tRNA, fMet-tRNA^iMet^ was purified with phenol/chloroform extraction, flash-frozen in liquid nitrogen stored in monosome buffer (20 mM HEPES-KOH pH 7.6, 100 mM KCl, 40 mM MgCl_2_, 1 mM DTT) at -80 °C. The efficiency of the reaction was assessed by RNA Mass Spectrometry.

### Reconstitution of mitochondrial initiation complexes *in vitro*

For the reconstitution of the pre-initiation complexes mtPIC-1 and mtPIC-2, 2.7 µg/µL mtSSU (OD280) were mixed with 500 nM rps3 mRNA (51-nucleotide sequence spanning the AUG codon, purchased from IDT) and 500nM mtIF3 and were incubated for 2 min at 37 °C, before being placed on ice for 10 min. For the formation of the pre-initiation complex mtPIC-3, 2.7 µg/µL mtSSU (OD280), 500 nM rps3 mRNA, and 500nM mtIF3 were mixed and incubated for 2 min at 37 °C. Then the initiator tRNA was introduced into the reaction as ternary complex in a 1:4 dilution. The ternary complex was formed by incubating 2.5 μM mtIF2 with 5 mM Guanosine 5′-[β,γ-imido]triphosphate trisodium salt hydrate (GDPNP, Sigma-Aldrich) for 30 min at room temperature before adding 2.5 μM fMet-tRNA^iMet^. After the addition of the ternary complex, the reaction was further incubated for 30 min at room temperature and for 10 min on ice. For the reconstitution of the initiation complex mtIC*, 3 µg/µL mtLSU (OD280) was added to the reaction, as an additional step to the described reaction of mtPIC-3, and incubated for another 3 min to induce monosome formation and incubated on ice for 10 min prior to grid preparation. All reconstitution reactions were carried out in Initiation buffer (20 mM HEPES-KOH, pH 7.6, 75 mM KCl, 40 mM MgCl_2_).

### Cryo-EM grid preparation

4 µL of the reconstitution reactions were applied onto Quantifoil R2/2 200 copper mesh grids, pre-coated with a 2 nm continuous carbon film and glow-discharged (3 mA for 20 sec). Samples were incubated on the grids for 20 sec and then blotted with filter paper for 3.5 sec in a temperature and humidity controlled Vitrobot Mark IV (T = 4°C, humidity 100%, blot force 2) followed by vitrification in liquid ethane.

### Single particle cryo-EM Data collection

Data collection was performed using a 300kV G4 Titan Krios electron microscope (Thermo Fisher) equipped with Falcon4i camera and a Selectris X energy filter using EPU for automated data acquisition at the Synchrotron SOLEIL. Data was collected at a nominal underfocus of – 0.4 to −2.2 µm at a magnification of 130,000x, yielding a pixel size of 0.94 Å. For mtPIC-1 and mtPIC-2, two datasets (1 and 2) were collected as movie stacks from two different grids, comprising 9,625 and 13,457 micrographs, respectively, and exported as EER files. Each movie stack was fractionated into 40 frames, corresponding to a total electron dose of 40 e^−^/Å^2^. For mtPIC-3, two datasets (3 and 4) were acquired as movie stacks from two separate grids, containing 20,008 and 26,051 micrographs, respectively, and exported as EER files. Each movie stack was fractionated into 40 frames, corresponding to a total electron dose of 40 e^−^/Å^2^. For mtIC*, dataset 5 was recorded as movie stacks with 34,059 micrographs and exported as EER files. Each movie stack was fractionated into 40 frames, corresponding to a total electron dose of 40 e^−^/Å^2^.

### Single particle cryo-EM Data processing

For all complexes, the complete processing pipeline was performed in cryoSPARC, with the workflows presented in Supplementary Fig. 4, 5, and 6. Cryo-EM validation parameters are provided in Supplementary Table 1.

For mtPIC-1 and mtPIC-2, pre-processing steps, particle picking as well as 2D classification and initial 3D classification were performed independently for datasets 1 and 2, and the resulting high-quality particles were subsequently merged. Gain and motion correction was performed using Patch motion correction and CTF estimation was carried out with Patch CTF. Following these steps, 9,508 and 12,272 micrographs for datasets 1 and 2, respectively, were kept. Particle picking was performed using blob and template picker resulting collectively in 1,607,019 and 2,733,504 particles, for datasets 1 and 2, respectively. Particles were extracted with a box size of 600 pixels downsampled to 300 pixels (pixel size of 1.88 Å), and were subjected to 2D classification, which retained 545,039 particles for dataset 1 and 888,747 for datasets 2. Particles were further filtered through 3D classification including Ab-initio, Heterogeneous refinement and 3D variability analysis, and duplicate particles were removed resulting in 316,853 and 496,866 particles for datasets 1 and 2, respectively which were combined. Particles were subsequently subjected to 3D classification with a mask on the SSU body, revealing a class of 420,241 particles containing mtIF3. These particles were re-extracted for high-resolution refinement using a 720-pixel box (0.94 Å/pixel). Global refinement yielded a resolution of 2.20 Å, which was further improved to 2.06 Å after Local and Global CTF refinement. Classes corresponding to pre-initiation complexes mtPIC-1 and mtPIC-2, were distinguished in a 3D classification with a focus mask on the exit site of the mRNA channel. This identified a class of 141,355 particles for mtPIC-1 with no mRNA present along the mRNA channel and a class of 62,215 particles for mtPIC-2 with mRNA present. Global refinement of the mtPIC-1 class resulted in a 2.28 Å resolution, whereas local refinements of the SSU head, SSU body and SSU foot using their respective focus masks reached 2.23 Å, 2.25 Å and 2.85 Å, respectively. Likewise, Global refinement of the mtPIC-2 class yielded a 2.55 Å resolution, while local refinements of the SSU head, SSU body and SSU foot achieved resolutions of 2.45 Å, 2.50 Å, and 3.14 Å.

For mtPIC-3, pre-processing, particle picking, 2D classification and initial 3D classification were carried out independently for datasets 3 and 4, and the selected, high-quality particles were subsequently combined. After Patch Motion Correction and Patch CTF, 19,820 and 25,617 micrographs for datasets 3 and 4, respectively, were accepted for further analysis. Particle picking was carried out using blob and template picker generating collectively 2,537,831 particles for dataset 3 and 4,020,330 particles for dataset 4. Particles were extracted using a 600-pixel box and downsampled to 300 pixels (1.88 Å/pixel). After 2D classification, 497,791 and 696,639 particles were selected for datasets 3 and 4, respectively. Additional particle cleaning was performed using 3D classification, as described above, and 316,412 particles for dataset 3 and 418,729 for datasets 4 were accepted. Particles were merged and passed through another round of 3D classification which produced a class of 330,546 particles containing mtIF3. The containing particles were re-extracted for high-resolution refinement using a 720-pixel box (0.94 Å/pixel). Global refinement reached a resolution of 2.64 Å, which was further enhanced to 2.40 Å through Local and Global CTF refinement. mtPIC-3 class was isolated through 3D classification using a focus mask on the mtIF3-tRNA area. The mtPIC-3 class comprised 18,894 particles, that reached a global refinement of 3.05 Å resolution, while local refinements of the SSU head and body achieved resolutions of 2.86 Å and 2.86 Å, respectively.

For mtIC*, 33,551 micrographs from the corresponding dataset 5, were retained for downstream analysis following preprocessing steps including gain and motion correction and CTF estimation. Particle picking was carried out using blob and template picker which generated collectively 4,371,444 particles that were subsequently extracted with a box size of 720 pixels downsampled to 360 pixels (pixel size of 1.88 Å) and subjected to 2D classification. 1,060,289 were accepted and passed through a 3D cleaning step leading to 721,567 high-quality particles. A second round of 3D classification was applied, separating SSU, LSU, and full mitoribosome particles. The latter comprised a class of 311,454 particles that was subjected to a 3D classification with a focus mask on the position of the P-site tRNA. A class of 25,802 full mitoribosome particles containing a P-site tRNA was identified. Particles were re-extracted for high-resolution refinement using a 756-pixel box (0.94 Å/pixel). After an initial global refinement step the particles were filtered through another round of 3D classification that led to the mtIC* class with 18,503 particles. Global refinement yielded a resolution of 3.18 Å, which was further improved to 2.92 Å after Local and Global CTF refinement. Local refinements of the SSU and LSU were performed using their respective focus masks achieving resolutions of 2.88 Å and 2.80 Å, respectively.

### Model building and refinement

For the mitochondrial pre-initiation complexes the recently published model of the cauliflower mitoribosome ^21^ (PDB: 9GYT) was used as a starting template and was fit into the respective high-resolution cryo-EM maps using rigid-body docking in ChimeraX ^28^. The corresponding AlphaFold2 ^29^ models for *Arabidopsis thaliana* mtIF3 (UNIPROT: Q6NLP2) and initiator tRNA were acquired and subsequently body-rigid fitted into their respective maps. The complete model for each complex was initially refined with restraints using the *phenix*.*real_space_refine* tool in PHENIX ^30^. The main components of each model were then automatically refined in PHENIX against their respective local refinement maps followed by manual refinement in COOT ^31^. Chimeric maps were generated using the Map Box option in PHENIX to extract density within 3Å of the model atoms. These maps were then fitted into the consensus maps in ChimeraX and merged using the *vop maximum* command. Model geometry was validated with MolProbity ^32^.

### Tissue Immunostaining of Arabidopsis roots

Roots of 10-day-old seedlings were fixed in PEM buffer (50 mM PIPES pH 6.9, 5 mM EGTA, 5 mM MgSO_4_) supplemented with 3.2% paraformaldehyde for 40 min and washed three times with PEM buffer for 5 min. The roots were digested in an enzymatic solution of PEM buffer supplemented with 2.5% (w/v) pectinase, 2.5% (w/v) cellulase, and 2.5% (w/v) pectolyase (Yakult Pharmaceutical) for 10 min. After removing the enzymatic solution, the roots were washed three times with PEM buffer for 5 min. Blocking of unspecific binding was achieved by incubation with PEM buffer supplemented with 8% (w/v) BSA and 0.1% Triton X-100 for 1 h at room temperature. After blocking, the roots were incubated with the primary antibodies anti-V5 (Thermo Fisher Scientific) and anti-HA (Cell Signaling Technology) overnight at 4 °C. The antibody solution was removed, and the tissue was washed five times with PEM buffer for 10 min. Then incubated with secondary antibodies anti-mouse Alexa 488 (Thermo Fisher Scientific) and anti-rabbit Alexa 568 (Thermo Fisher Scientific) overnight at 4 °C. The antibody solution was removed, and the tissue was washed five times with PEM buffer for 10 min. The roots were counter-stained with DAPI and mounted on a microscope slide with a drop of antifade medium (Vectashield). All antibodies were diluted in their working concentration in PEM buffer supplemented with 1% (w/v) BSA.

### Confocal imaging

Confocal imaging was performed with a Zeiss LSM 980 with Airyscan 2 inverted microscope with a 63x oil-immersion objective or a Leica SP8 inverted microscope with a 63x oil-immersion objective. The following combinations of laser excitation and emission filters were used for various fluorophores: DAPI (405 nm laser excitation, 445/40 nm emission), AlexaFluor 488 (491 nm laser excitation, 528/38 nm emission) and AlexaFluor 568 (561 nm laser excitation, 617/73 nm emission).

### Figure preparation

Figures were prepared using BioRender (biorender.com) and ChimeraX, developed by the Resource for Biocomputing, Visualization, and Informatics at the University of California, San Francisco, with support from National Institutes of Health R01-GM129325 and the Office of Cyber Infrastructure and Computational Biology, National Institute of Allergy and Infectious Diseases ^33^.

## Supplementary Figures

**Supplementary Figure 1:** Subcellular localization and production of the plant mitochondrial initiation factors 2 and 3.

**a** Characterization of plant mitochondrial initiation factors localizations. Tissue immunostaining of Arabidopsis roots expressing mtIF2 and mtIF3. Expression of mtIF2 is detected by anti-V5/alexa488, whereas mtIF3 is detected by anti-HA/alexa568. Nuclei are stained with DAPI. The overlay channel highlights the co-localization of initiation factors in mitochondria, indicated by white arrows. Scale bars represent 10 μm. **b** Expression and affinity purification of initiation factors in tobacco cell extract. SDS-PAGE showing the total protein content after expression of mtIF2 and mtIF3 (lysate), the elution steps (elution), and the concentrated proteins after removal of the affinity tags (amicon). Green arrows indicate mtIF2, while red arrows indicate mtIF3. Molecular weight markers are shown on the left of the respective protein blots.

**Supplementary Figure 2:** Sequence alignment and structural comparison of (mt)IF3 from

*Arabidopsis thaliana, Homo sapiens*, and *Thermus thermophilus*. Comparative analysis of mtIF3. **a** Multiple amino acid sequence alignment of the three (mt)IF3 orthologs generated using Clustal Omega, highlighting conserved residues and main domains. **b** Superposition of the plant mtIF3/fMet-tRNA^iMet^ complex (mtPIC-3) presented here with *H. sapiens* mtIF3 (PDB: 6RW4) and *T. thermophilus* IF3 (PDB: 5LMQ).

**Supplementary Figure 3:** Preparation and characterization of plant mitochondria initiator tRNA *In vitro* transcription, acylation and formylation of the plant initiator tRNA. **a** *In vitro* transcription of Arabidopsis mitochondria initiator tRNA, with molecular weight markers shown on the left. **b** Schematic representation of the aminoacylation and formylation reactions of the initiator tRNA. MetRS first attaches methionine to the 3′ end of the initiator tRNA, after which FT adds a formyl group to the charged methionine. The scheme was created in BioRender (biorender.com). **c** SDS–PAGE analysis of the main fractions, including the lysate (Lys), soluble fraction (Sol), flow-through (Flow), and eluate from IMAC purification of the MetRS and FT enzymes, boxed in black and red dashed lines respectively. Molecular weight markers shown on the left. **d** RNA mass spectrometry chromatogram characterizing the result of the aminoacylation and formylation reactions of the initiator tRNA. Peaks corresponding to uncharged tRNA molecules and successfully aminoacylated and formylated tRNA molecules are shown in black and magenta respectively.

**Supplementary Figure 4:** Single-particle data processing workflow of the mitochondrial pre-initiation complexes mtPIC-1 and mtPIC-2.

Schematic overview of the data processing workflow. **a** Pre-processing steps followed by 2D and 3D classification leading to global refinement of mtPIC-1 and mtPIC-2. An orientation distribution plot is shown for the globally refined maps. **b** Local refinements of mtPIC-1 and mtPIC-2, with all masks used indicated. For the final reconstruction, Gold-standard Fourier shell correlation (GSFSC) plots are shown, with resolution determined at the 0.143 threshold. Local resolution maps are displayed on a consistent resolution scale, shown in both front-view and cut-view representations.

**Supplementary Figure 5:** Single-particle data processing workflow of the mitochondrial pre-initiation complex mtPIC-3.

Schematic overview of the data processing workflow. **a** Pre-processing steps followed by 2D and 3D classification leading to global refinement of mtPIC-3. An orientation distribution plot is shown for the globally refined map. **b** Local refinements of mtPIC-3, with all masks used indicated. For the final reconstruction, Gold-standard Fourier shell correlation (GSFSC) plots are shown, with resolution determined at the 0.143 threshold. Local resolution maps are displayed on a consistent resolution scale, shown in both front-view and cut-view representations.

**Supplementary Figure 6:** Single-particle data processing workflow of the mitochondrial initiation complex mtIC*.

Schematic overview of the data processing workflow. **a** Pre-processing steps followed by 2D and 3D classification leading to global refinement of mtIC*. An orientation distribution plot is shown for the globally refined map. **b** Local refinements of mtIC*, with all masks used indicated. For the final reconstruction, Gold-standard Fourier shell correlation (GSFSC) plots are shown, with resolution determined at the 0.143 threshold. Local resolution maps are displayed on a consistent resolution scale, shown in both front-view and cut-view representations.

## Supplementary Table

**Supplementary Table 1:** Cryo-EM data collection, refinement and validation statistics.

## Data availability

The single particle cryo-EM maps of *B. oleracea* translation initiation complexes have been deposited at the Electron Microscopy Data Bank (EMDB) and models on the protein data bank (PDB). Plant mitochondrial pre-initiation complex 1 (mtPIC-1) PDB XXXX. Plant mitochondrial pre-initiation complex 2 (mtPIC-2) PDB XXXX. Plant mitochondrial pre-initiation complex 3 (mtPIC-3) PDB XXXX. Plant mitochondrial initiation complex (mtIC*) PDB XXXX.

## Acknowledgements

We thank Florent Waltz (Biozentrum, University of Basel) for advice on cryo-EM data analysis and presentation; Nicolas Baumberger (Protein Production and Purification platform, Institut de Biologie Moléculaire des Plantes, CNRS) for advice on protein purification; and Philippe Wolff (Institut de Biologie Moléculaire et Cellulaire, CNRS) for validation of tRNA acylation by RNA mass spectrometry.

This work was supported by the “Centre National de la Recherche Scientifique”, the University of Strasbourg, by Agence Nationale de la Recherche (ANR) grants ‘DAMIA, ANR-20-CE11-0021’ and ‘PROPHAN, ANR-22-CE12-0008-01’ to PG and by the LabEx consortium “MitoCross” in the frame of the French National Program “Investissement d’Avenir” ANR-11-LABX-0057_MITOCROSS. We acknowledge SOLEIL and the French EQUIPEX+ France Cryo-EM (ANR-21-ESRE-0046) for provision of cryo-EM facilities. This work benefited from the cryo-EM platform of I2BC supported by the French Infrastructure for Integrated Structural Biology (FRISBI) [ANR-10-INSB-05-05].

## Author contributions

VS performed all experiments. HS performed cryo-EM sample preparation and data acquisition. VS and HS performed cryo-EM data processing, model building and interpretation. VS and PG conceived the project and wrote the manuscript.

## Competing interests

The authors declare no competing interests.

## Notes

### Competing Interest Statement

The authors have declared no competing interest.

