## Supplementary Figure 1 for "Missing steps of mitochondrial translation initiation identified in plants"

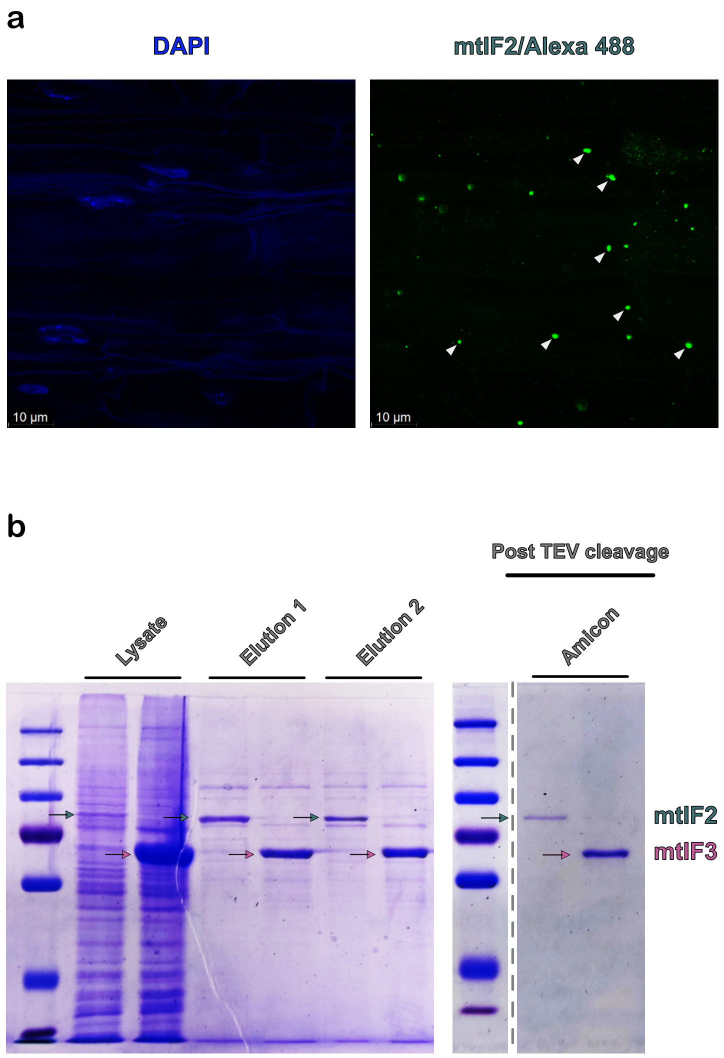


**Supplementary Figure 1**: Subcellular localization and production of the plant mitochondrial initiation factors 2 and 3.

**a** Characterization of plant mitochondrial initiation factors localizations. Tissue immunostaining of Arabidopsis roots expressing mtIF2 and mtIF3. Expression of mtIF2 is detected by anti-V5/alexa488, whereas mtIF3 is detected by anti-HA/alexa568. Nuclei are stained with DAPI. The overlay channel highlights the co-localization of initiation factors in mitochondria, indicated by white arrows. Scale bars represent 10 µm. **b** Expression and affinity purification of initiation factors in tobacco cell extract. SDS-PAGE showing the total protein content after expression of mtIF2 and mtIF3 (lysate), the elution steps (elution), and the concentrated proteins after removal of the affinity tags (amicon). Green arrows indicate mtIF2, while red arrows indicate mtIF3. Molecular weight markers are shown on the left of the respective protein blots.
