## Supplementary Figure 2 for "Missing steps of mitochondrial translation initiation identified in plants"

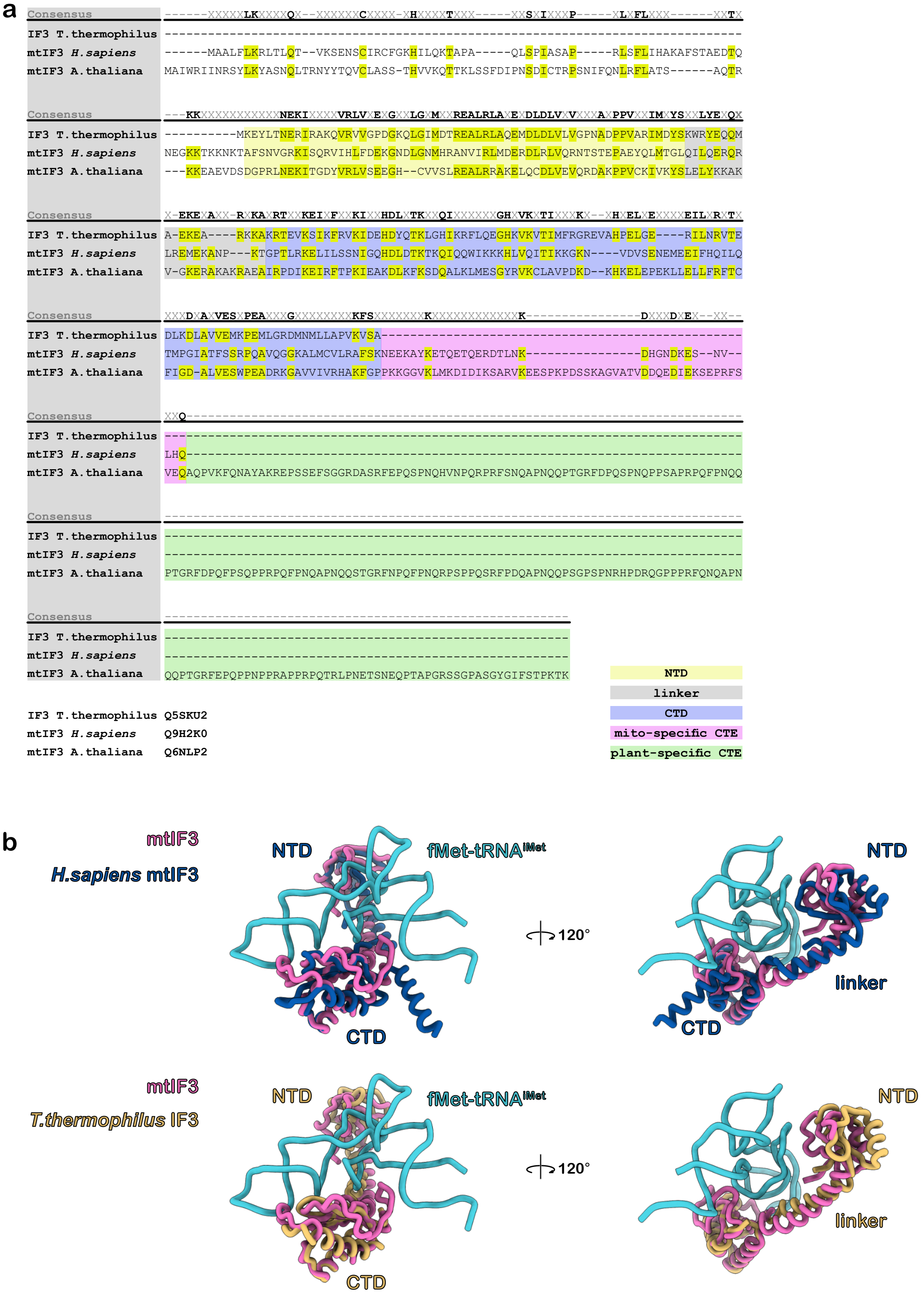


**Supplementary Figure 2**: Sequence alignment and structural comparison of (mt)IF3 from *Arabidopsis thaliana*, *Homo sapiens*, and *Thermus thermophilus*.

Comparative analysis of mtIF3. **a** Multiple amino acid sequence alignment of the three (mt)IF3 orthologs generated using Clustal Omega, highlighting conserved residues and main domains. **b** Superposition of the plant mtIF3/fMet-tRNA$^{iMet}$ complex (mtPIC-3) presented here with *H. sapiens* mtIF3 (PDB: 6RW4) and *T. thermophilus* IF3 (PDB: 5LMQ).
