## Supplementary Figure 3 for "Missing steps of mitochondrial translation initiation identified in plants"

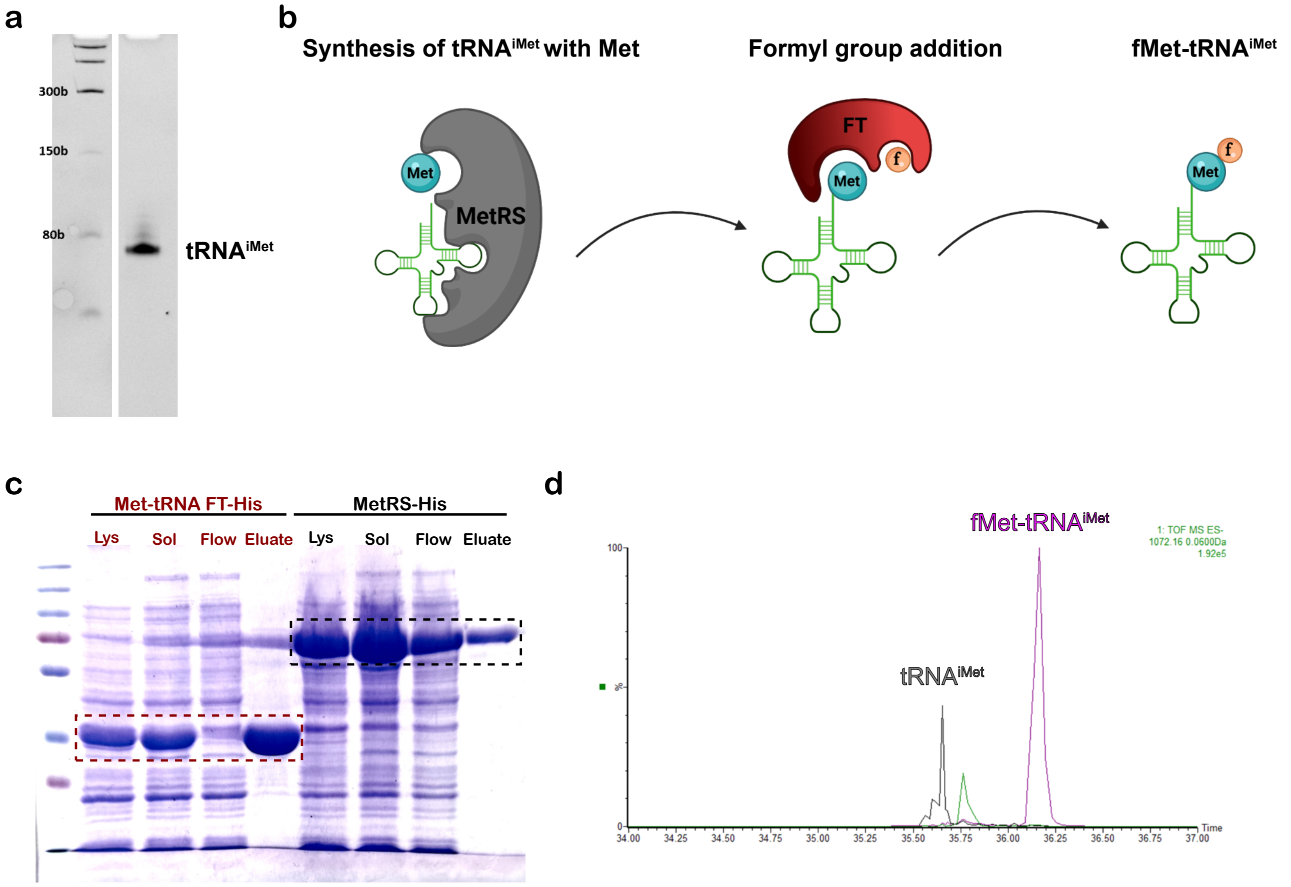


**Supplementary Figure 3**: Preparation and characterization of plant mitochondria initiator tRNA

*In vitro* transcription, acylation and formylation of the plant initiator tRNA. **a** *In vitro* transcription of Arabidopsis mitochondria initiator tRNA, with molecular weight markers shown on the left. **b** Schematic representation of the aminoacylation and formylation reactions of the initiator tRNA. MetRS first attaches methionine to the 3′ end of the initiator tRNA, after which FT adds a formyl group to the charged methionine. The scheme was created in BioRender (biorender.com). **c** SDS–PAGE analysis of the main fractions, including the lysate (Lys), soluble fraction (Sol), flow-through (Flow), and eluate from IMAC purification of the MetRS and FT enzymes, boxed in black and red dashed lines respectively. Molecular weight markers shown on the left. **d** RNA mass spectrometry chromatogram characterizing the result of the aminoacylation and formylation reactions of the initiator tRNA. Peaks corresponding to uncharged tRNA molecules and successfully aminoacylated and formylated tRNA molecules are shown in black and magenta respectively.
