## Supplementary Figure 6 for "Missing steps of mitochondrial translation initiation identified in plants"

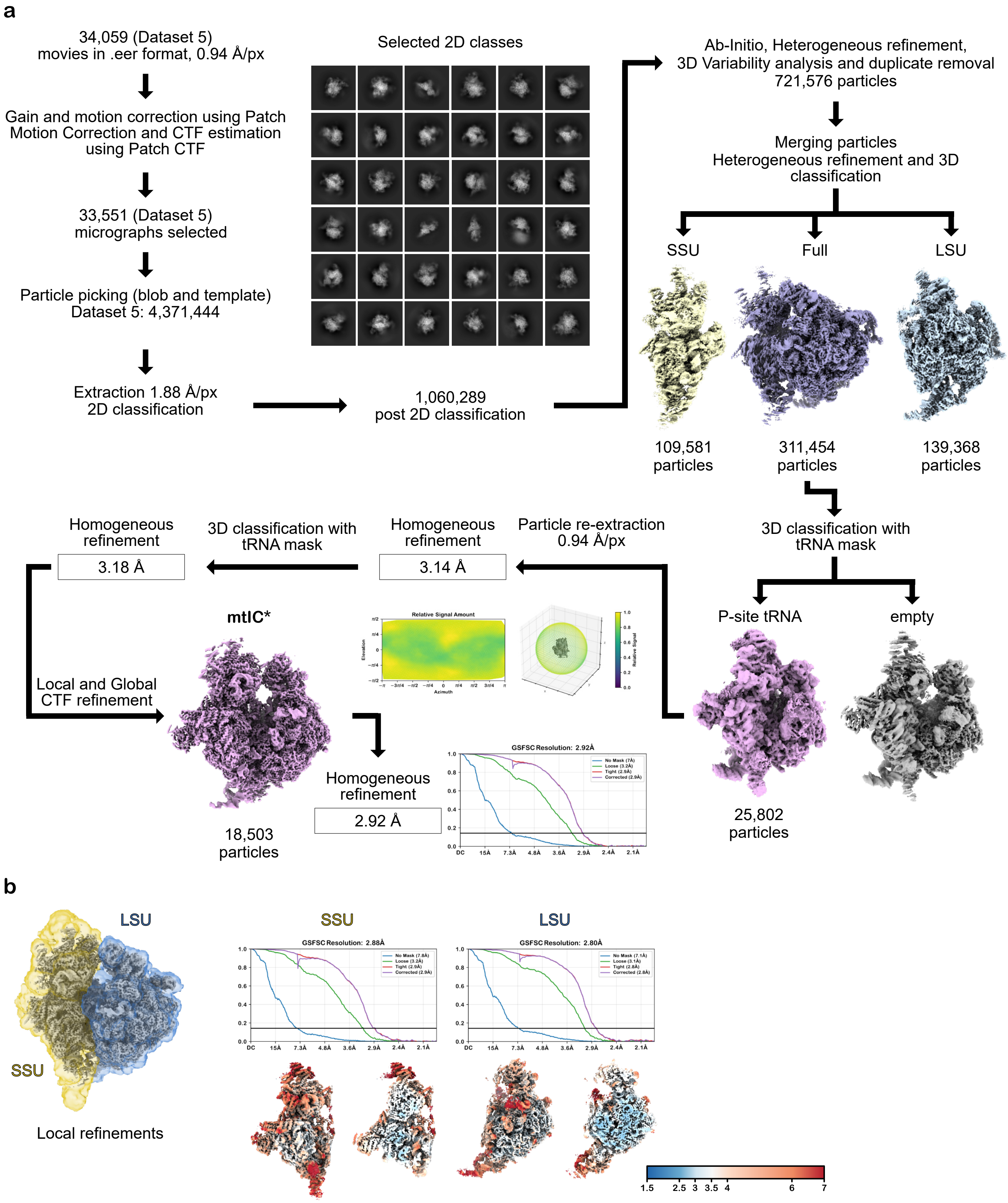


**Supplementary Figure 6**: Single-particle data processing workflow of the mitochondrial initiation complex mtIC*

Schematic overview of the data processing workflow. **a** Pre-processing steps followed by 2D and 3D classification leading to global refinement of mtIC*. An orientation distribution plot is shown for the globally refined map. **b** Local refinements of mtIC*, with all masks used indicated. For the final reconstruction, Gold-standard Fourier shell correlation (GSFSC) plots are shown, with resolution determined at the 0.143 threshold. Local resolution maps are displayed on a consistent resolution scale, shown in both front-view and cut-view representations.
