## Supplementary Table 1 for "Missing steps of mitochondrial translation initiation identified in plants"

| | mtPIC-1 | mtPIC-2 | mtPIC-3 | mtIC-1* |
|---|---|---|---|---|
| **PDB** | **XXXX** | **XXXX** | **XXXX** | **XXXX** |
| **Unfocused** | **XXXX** | **XXXX** | **XXXX** | **XXXX** |
| **Data collection and processing** | | | | |
| Magnification | 130,000x | 130,000x | 130,000x | 130,000x |
| Voltage (kV) | 300 | 300 | 300 | 300 |
| Electron exposure ($e^-/Å^2$) | 40 | 40 | 40 | 40 |
| Defocus range (μm) | -0.4 to -2.2 | -0.4 to -2.2 | -0.4 to -2.2 | -0.4 to -2.2 |
| Pixel size (Å) | 0.94 | 0.94 | 0.94 | 0.94 |
| Symmetry imposed | C1 | C1 | C1 | C1 |
| Initial particle images (no.) | 4,340,523 | 4,340,523 | 6,558,161 | 4,371,444 |
| Final particle images (no.) | 141, 355 | 62, 215 | 18, 894 | 18, 503 |
| Map resolution (Å) FSC 0.143 | 2.28 | 2.55 | 3.05 | 2.92 |
| Map resolution range (Å) | 2.2 - 7.3* | 2.1 - 8.2* | 2.1 - 10.6* | 2.1 - 9.3* |
| **Refinement** | | | | |
| Initial model used (PDB code) | 9GYT | 9GYT | 9GYT | 9EVS |
| Model resolution (Å) FSC 0.143 | 2.3 | 2.55 | 3.05 | 2.92 |
| CC Model vs Data (mask) | 0.78 | 0.78 | 0.79 | 0.78 |
| Map sharpening *B* factor ($Å^2$) | -30.8 | -28.4 | -12.9 | -22.2 |
| Model composition | | | | |
| Non-hydrogen atoms | 89548 | 89124 | 92977 | 207751 |
| Residues : Protein - Nucleotide | 6707 - 1610 | 6632 - 1619 | 6863 - 1681 | 13343 - 4686 |
| Ligands | ATP: 1 | ATP: 1 | ATP: 1 | ATP: 1 |
| *B* factors ($Å^2$) | | | | |
| Protein | 0.00/235.67/47.95 | 0.00/235.67/48.35 | 0.00/540.51/75.72 | 0.00/444.13/54.33 |
| Nucleotide | 1.55/271.67/54.66 | 1.55/272.81/54.72 | 8.32/562.91/71.13 | 0.37/376.89/66.33 |
| Ligand | 12.07/76.37/30.71 | 12.07/76.37/30.71 | 3.00/95.27/31.17 | 4.98/97.19/34.70 |
| R.m.s. deviations | | | | |
| Bond length ($Å^2$) | 0.007 | 0.007 | 0.007 | 0.004 |
| Bond angles (°) | 1.000 | 0.998 | 0.945 | 0.811 |
| Validation | | | | |
| MolProbity score | 2.06 | 2.09 | 2.24 | 1.90 |
| Clash score | 11.86 | 12.07 | 16.19 | 6.46 |
| Poor rotamers (%) | 2.20 | 2.15 | 2.23 | 2.71 |
| Ramachandran plot | | | | |
| Favored (%) | 96.64 | 96.40 | 96.07 | 96.60 |
| Allowed (%) | 3.18 | 3.41 | 3.74 | 3.34 |
| Disallowed (%) | 0.18 | 0.18 | 0.19 | 0.06 |

* min - 75th percentile, in cryoSPARC Local resolution Estimation

**Extended Data Table 1:** Cryo-EM data collection, refinement and validation statistics
